# Detection of arterial wall abnormalities via Bayesian model selection

**DOI:** 10.1101/422485

**Authors:** Karen Larson, Clark Bowman, Costas Papadimitriou, Petros Koumoutsakos, Anastasios Matzavinos

## Abstract

Patient-specific modeling of hemodynamics in arterial networks has so far relied on parameter estimation for inexpensive or small-scale models. We describe here a Bayesian uncertainty quantification framework which makes two major advances: an efficient parallel implementation, allowing parameter estimation for more complex forward models, and a system for practical model selection, allowing evidence-based comparison between distinct physical models. We demonstrate the proposed methodology by generating simulated noisy flow velocity data from a branching arterial tree model in which a structural defect is introduced at an unknown location; our approach is shown to accurately locate the abnormality and estimate its physical properties even in the presence of significant observational and systemic error. As the method readily admits real data, it shows great potential in patient-specific parameter fitting for hemodynamical flow models.

## 1 Introduction

Mathematical models for hemodynamics trace back to the work of Euler, who described a one-dimensional treatment of blood flow through an arterial network with rigid tubes [11, 33]; more sophisticated one-dimensional models are still used to study a variety of physio-pathological phenomena [1, 2, 13, 21, 23, 29, 30, 37]. Computational advances have also allowed for the development of computationally intensive three-dimensional models [12, 14, 32, 34, 39, 40], which have been used to accurately simulate specific human arteries (e.g., the carotid arteries [18]) and model their material properties (e.g., of cerebral arterial walls [38]). There also exist multicomponent models [10], which are amenable to applications such as modeling oxygen transport to solid tumors [6] and surgical tissue flaps [24, 25].

Despite the sophistication of these approaches, there remain a number of challenges in the creation of patient-specific models using individual medical data. In particular, computational expense usually limits arterial parameter estimation to the one-dimensional class of models [33, 14], which have nonetheless proven sufficiently robust to study fluid-structure interactions and viscoelasticity [30, 23] and create a patient-specific model for vascular bypass surgery [20]. Several approaches exist for parameter estimation and uncertainty quantification for these models. Gradient descent has been used to estimate arterial compliance parameters [22], recovering single parameters assumed constant in space and time. Sensitivity analysis has also been used, successfully quantifying output sensitivity to various uncertainties in a stochastic flow network [7]. More recently, computational methods based upon Bayesian optimization and multi-fidelity information fusion for model inversion have been explored [31].

The chief contribution of this work is to introduce a Bayesian framework for uncertainty quantification in a bifurcating network of one-dimensional extensible arteries. The advantages of the approach are twofold. First, it utilizes transitional Markov chain Monte Carlo (TMCMC), a highly parallelizable algorithm for approximate sampling which allows practical uncertainty quantification even for large arterial networks; our high-performance implementation Π4U will be shown to simultaneously and efficiently estimate several unknown parameters in this setting. Second, the approach can practically be used for Bayesian model selection, allowing for evidence-based comparison between models with distinct physical assumptions. The approach thus represents a significant advance in fitting patient-specific hemodynamical flow models.

Specifically, we consider a branching network of 19 arteries in which a structural flaw (e.g., an aneurysm) has been introduced at an unknown location. Sections 2 and 3 describe the onedimensional blood flow model and the uncertainty quantification framework. In Section 4, we use the flawed model to simulate noisy observations of the flow velocity at fixed points in the network. We then use Bayesian model selection to probabilistically locate the defect within the network and accurately recover its structural properties, showing the approach to be effective even when the model is misspecified (i.e., the model used to generate the artificial data differs from the model used to perform uncertainty quantification). As the method readily admits clinical blood flow data, which have been shown to be measurable with non-invasive procedures [36, 28, 35, 26], it shows great potential in diagnosing patient-specific structural issues in the circulatory system.

## 2 Nonlinear One-Dimensional Blood Flow Model

We first introduce the one-dimensional blood flow model. While such models can be derived via a scaling of the Navier-Stokes equations for viscous flow [27], we use here the geometry- and conservation-motivated approach described by Sherwin et al. [33] and Peiró and Venziani [14]. In this approach, the viscous, incompressible flow is assumed to move only in the axial direction (i.e., along the one-dimensional artery), to exhibit axial symmetry, and to maintain constant internal pressure over orthogonal cross-sections. The artery is assumed to have low curvature and to be distensible in the radial direction. A schematic of the artery appears in Fig. 1.

**Figure 1:**
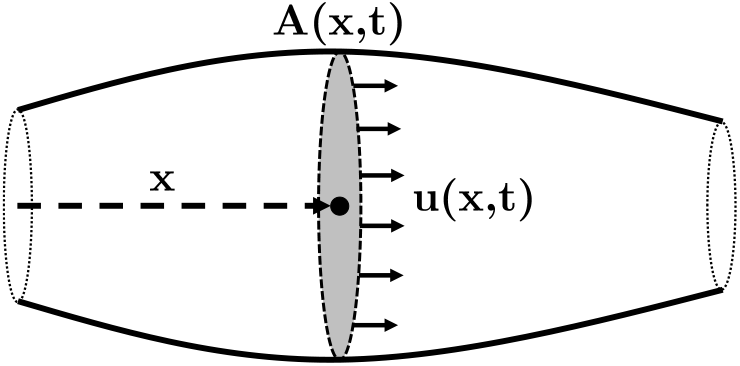
Schematic of one-dimensional artery segment.

The artery of constant length *ℓ* and position-dependent cross-sectional area *A*(*x*, *t*) is filled with blood flowing at velocity *u*(*x*, *t*) and with internal cross-sectional pressure *p*(*x*, *t*), yielding the crosssectional flux *Q*(*x*, *t*) = *A*(*x*, *t*)*u*(*x*, *t*). Choosing *u*, *A*, and *p* as the independent variables, the partial differential equation governing the incompressible flow can be derived from conservation of mass and momentum:

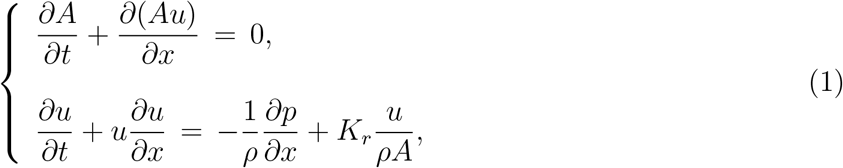

where *ρ* is the flow density and *K_r_* is a parameter representing viscous resistance per unit length, here given by 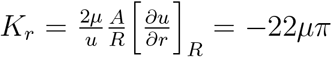 in terms of the channel radius *R*(*x*, *t*) and the viscosity *μ* of blood [30, 29].

The system is closed using a constitutive law to relate pressure and area. Using the Laplace tube law and assuming that the arterial wall is purely elastic,

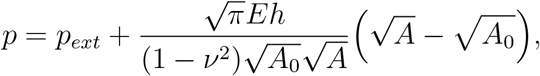

where *p_ext_* is the external pressure, *E* is the Young’s modulus of the wall, *h* is the wall thickness, *A*_0_ is the relaxed cross-sectional area, and *ν* is the Poisson ratio, here taken to be 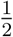. For notational simplicity, we collect the coefficient into a single stiffness parameter *B*, yielding

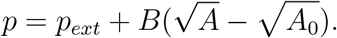

(1) can then be re-written in the form of a nonlinear hyperbolic conservation law [33]:

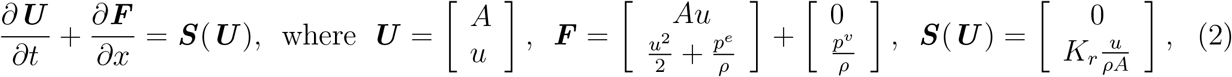

in terms of the elastic component *p^e^*(*x*, *t*) and viscoelastic component *p^v^*(*x*, *t*) of the pressure.

The hyperbolic system is approximated numerically using a discontinuous Galerkin method. The one-dimensional domain Ω = (*a*, *b*) is discretized into *N* non-overlapping elements 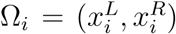 such that 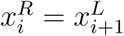 and 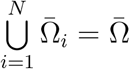; discrete approximations to the corresponding weak formulation are found in terms of orthonormal Legendre polynomials of degree *p* [9, 16] (see, e.g., [9, 17] for the advantages of this approach). Inlet and outlet boundary elements use upwind flux, while a second-order Adams-Bashforth scheme [9] is used for time integration.

To extend the model to a branching arterial network, multiple arteries are joined via coupled boundary conditions at bifurcations. An example of such a bifurcation appears in Figure 2. Boundary conditions are physically motivated; mass should be conserved through bifurcations, while momentum should be continuous at the boundary, i.e.,

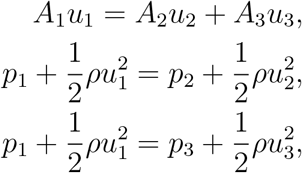

where *p_i_*, *A_i_*, and *u_i_* correspond to the ith artery. Other branching configurations appear in [33].

**Figure 2:**
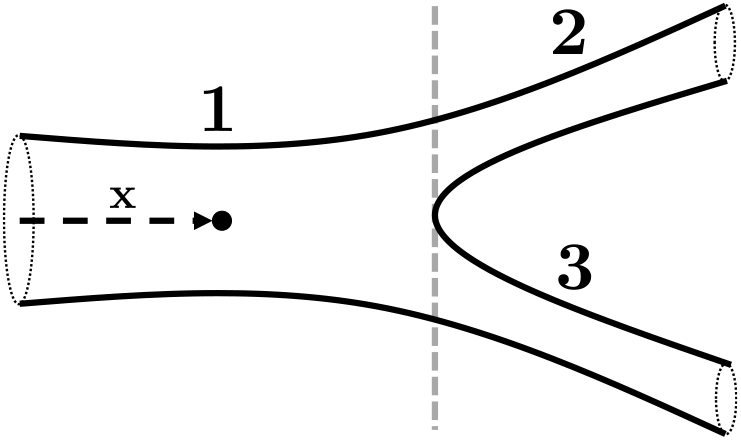
Schematic of Y-bifurcation in an arterial network.

## 3 Bayesian Uncertainty Quantification

The primary practical goal of this paper is to identify structural defects in an arterial network using observations of the blood flow velocity. By varying the material properties of the arteries, perturbations to the flow can be computed via the blood flow model described in Section 2; in this sense, the goal is to solve the inverse problem of determining structural parameters given velocity data as model output. In real applications, these velocity data may be corrupted by noise (e.g., measurement error). Furthermore, the model itself may be misspecified; for example, model parameters assumed as known may be incorrect. In this section, we introduce our recent Bayesian framework for uncertainty quantification which is amenable to the former issue (noise) and will prove robust to the latter (misspecification). Section 3.3 describes the TMCMC method which forms the core of this approach: its parallelizability allows for feasible application to more expensive models, such as the model of Section 2, via the use of high-performance computing.

### 3.1 Parameter Estimation

Denote as *M* the mathematical model of interest, which deterministically maps a set of *n* parameters 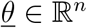 to *m* outputs 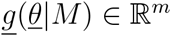 (here, <u>*g*</u> denotes the forward problem, and so <u>*g*</u>(·|*M*) is a solution to the forward problem using the model *M*). The inverse problem is then to estimate the parameters <u>*θ*</u> given the model outputs. We assume that these model outputs have been corrupted by noise (due to, e.g., measurement, computational, or modeling error) as

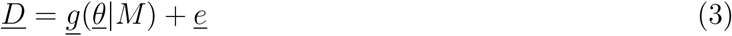

in terms of a random predictive error <u>*e*</u>. Under the Bayesian formulation of this problem, the parameters <u>*θ*</u> are assigned a prior distribution *π*(*<u>θ</u>*|*M*) given any *a priori* knowledge of the parameters based on, e.g., physical constraints; the posterior *p*(<u>*θ*</u>|<u>*D*</u>, *M*) that observed data <u>*D*</u> were generated by parameters <u>*θ*</u> can then be found as

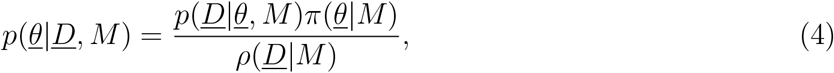

using the likelihood <u>*p*</u>(<u>*D*</u>|*<u>θ</u>*, *M*), calculated by evaluating <u>*g*</u>(<u>*θ*</u>|*M*) and using the form of <u>*e*</u>, and the evidence *ρ*(*<u>D</u>*|*M*) of the model class, computed via the multi-dimensional integral

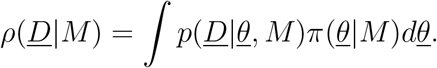

In order to calculate the likelihood *p*(*<u>D</u>*|*<u>θ</u>*, *M*), we make the simplifying assumption that <u>*e*</u> is normally distributed with zero mean and covariance matrix Σ, which may itself include additional unknown parameters. Since the model outputs <u>*g*</u> are deterministic, it follows that <u>*D*</u> is also normally distributed, and so the explicit likelihood *p*(*<u>D</u>*|*<u>θ</u>*, *M*) is given by

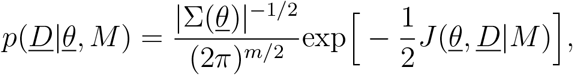

where

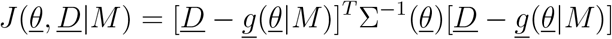

is the weighted measure of fit between the model predictions and the measured data, | · | denotes determinant, and the parameter set <u>*θ*</u> is augmented to include parameters that are involved in the structure of the covariance matrix Σ.

### 3.2 Model Selection

The Bayesian approach to uncertainty quantification is especially useful in the context of model selection. The evidence *ρ*(<u>*D*</u>|*M*) which appears in (4) is a measure of the degree to which the model M can explain the data *<u>D</u>*; when *M* is one particular model in a parameterized class 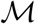 of models, the evidence can be used to derive a distribution on models. Let *Pr*(*M_i_*) be a prior distribution on models in the class 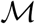. The posterior *Pr*(*M_i_*|<u>*D*</u>) can again be derived from Bayes’ theorem:

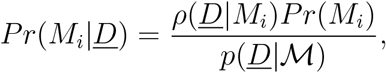

where 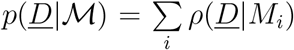 is a normalization constant. Intuitively, *Pr*(*M_i_*|<u>*D*</u>) is a distribution which describes the probability of the data <u>*D*</u> having been generated from model *M_i_* (as opposed to another model *M_j_*) under the assumption that at least one model in 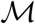 is the true model, i.e., was actually used to generate the data. If a uniform prior is assumed on models, this posterior is directly proportional to the evidence *ρ*(*<u>D</u>*|*M_i_*), and so model selection is “free” when the evidence is already calculated for parameter estimation [4, 15, 5, 19].

### 3.3 Transitional Markov Chain Monte Carlo

The main computational barrier in calculating the posterior distribution (4) is the complex forward problem <u>*g*</u> (here, the blood flow model of Section 2) which appears in the fitness *J*(<u>*θ*</u>, *<u>D</u>*|*M*). Our specific implementation Π4U of Bayesian uncertainty quantification has two advantages in this respect: first, it approximately samples the posterior via transitional Markov chain Monte Carlo (TMCMC), which is massively parallelizable, and second, it leverages an efficient parallel architecture for task sharing described in Appendix A.

The TMCMC algorithm used functions by smoothly transitioning to the target distribution (the posterior *p*(*<u>θ</u>*|*<u>D</u>*, *M*)) from the prior *π*(*<u>θ</u>*|*M*). To accomplish this, a series of intermediate distributions are constructed iteratively:

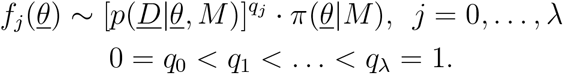

The explicit algorithm is summarized below in Algorithm 1. It begins by taking *N*_0_ samples <u>*θ*</u>_0,*k*_ from the prior distribution *f*_0_(<u>*θ*</u>) = *π*(*<u>θ</u>*|*M*). For each stage *j* of the algorithm, the current samples are used to compute the plausibility weights *w*(*<u>θ</u>_j,k_*) as

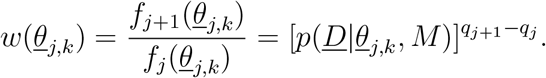

Recent literature suggests that *q*_*j*+1_, which determines how smoothly the intermediate distributions transition to the posterior, should be taken to make the covariance of the plausibility weights at stage *j* smaller than a tolerance covariance value, often 1.0 [8, 15].

**Algorithm 1.**
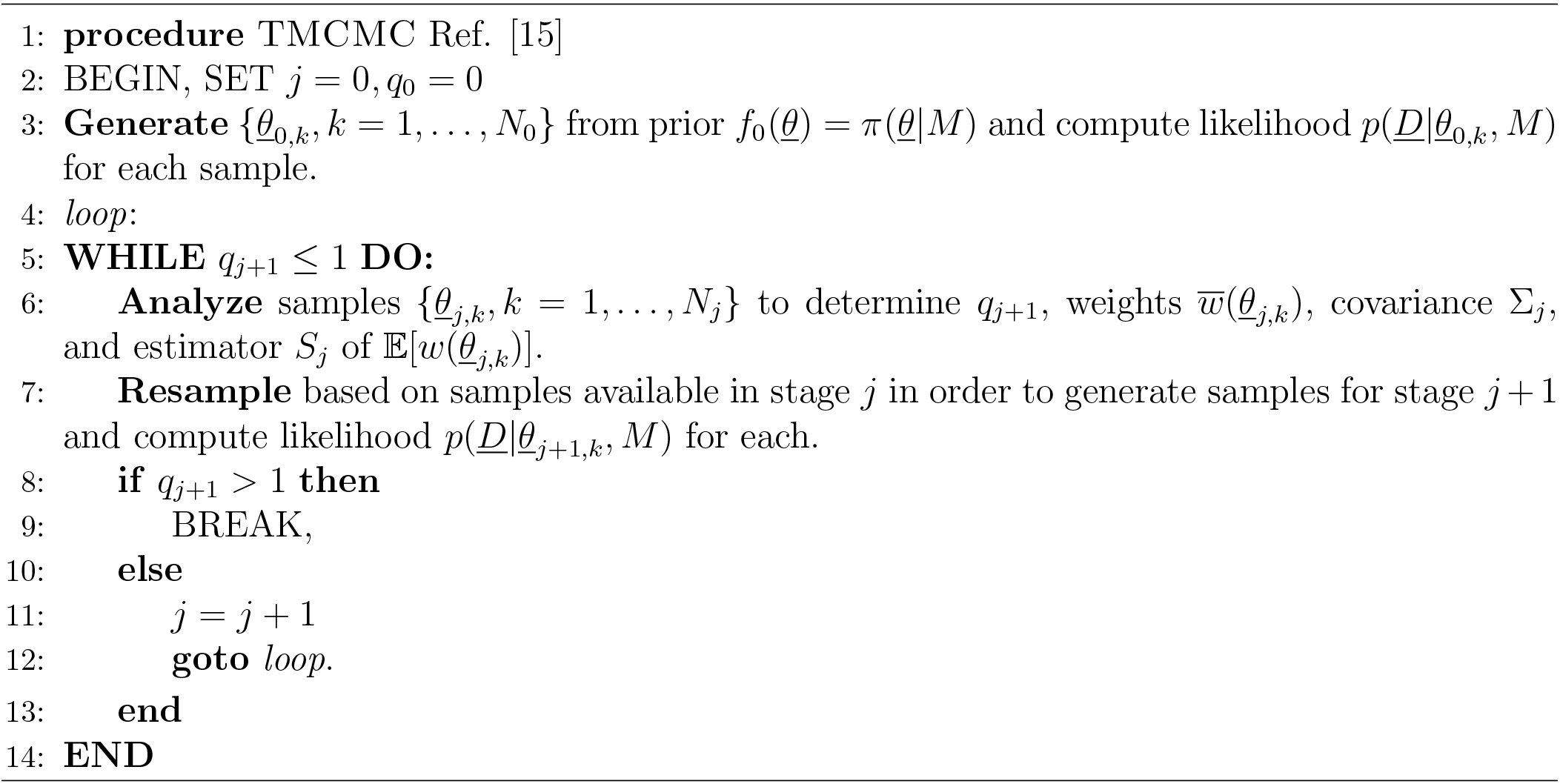
TMCMC.

Next, the algorithm calculates the average *S_j_* of the plausibility weights, the normalized plausibility weights 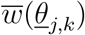, and the scaled covariance Σ*_j_* of the samples *<u>θ</u>_j,k_*, which is used to produce the next generation of samples *<u>θ</u>*_*j*+1,*k*_:

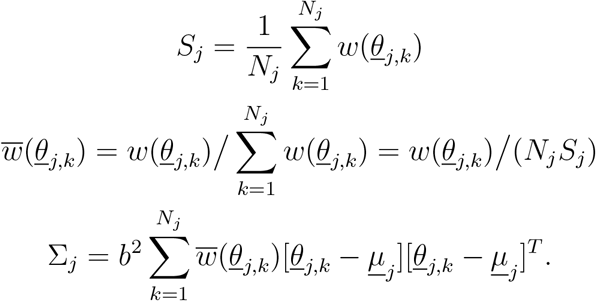

Σ*_j_* is calculated using the sample mean *<u>μ</u>_j_* and a scaling factor *b*, usually 0.2 [8, 15].

The algorithm then generates *N*_*j*+1_ samples 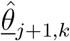 by randomly selecting from the previous generation {*<u>θ</u>_j, k_*} such that 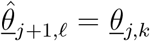 with probability 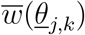. These samples are selected independently at random, so any parameter can be selected multiple times – call *n*_*j*+1,*k*_ the number of times *<u>θ</u>_j,k_* is selected. Each unique sample is used as the starting point of an independent Markov chain of length *n*_*j*+1,*k*_ generated using the Metropolis algorithm with target distribution and a Gaussian proposal distribution with covariance Σ*_j_* centered at the current value.

Finally, the samples *<u>θ</u>*_*j*+1,*k*_ are generated for the Markov chains, with *n*_*j*+1,*k*_ samples drawn from the chain starting at *<u>θ</u>_j,k_*, yielding *N*_*j*+1_ total samples. The algorithm then either moves forward to generation *j* + 1 or terminates if *q*_*j*+1_ > 1.

## 4 Results

We now apply the Bayesian framework of Section 3 to the blood flow model of Section 2. In particular, we study the example 19-artery network shown in Figure 3. The solution for our deterministic model, given by (2) and solved using a discontinuous Galerkin method with time step Δ*t*_1_ = 0.00004 seconds, plays the role of *<u>g</u>* in the model prediction equation (3). Measurements of the flow velocity are taken at *N* specified locations which vary by experiment and occur with a sampling period of Δ*t*_2_ = 1600Δ*t*_1_ = 0.064 seconds.

**Figure 3:**
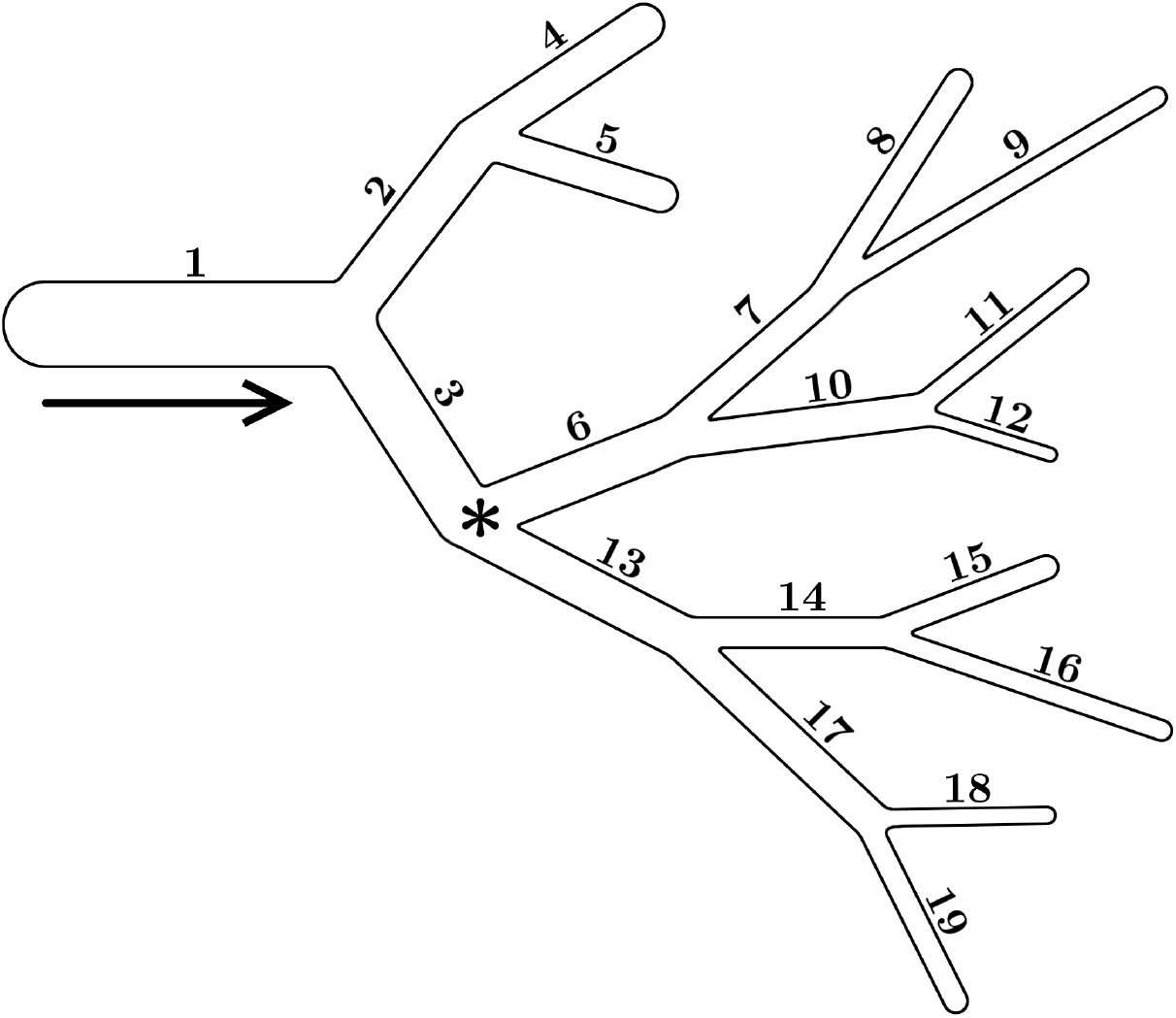
Schematic of arterial network (not to scale). 19 arteries have varied lengths (ranging from 0.026 to 0.17 *m*) and cross-sectional areas (ranging from 10^−5^ to 10^−6^ *m*^2^). The star shows an example measurement location at a bifurcation.

Blood (viscosity *μ*\* = 0.0045, with asterisks denoting reference values) in the network begins with zero velocity and is driven by a specified inflow velocity at the beginning of the first artery: a sum of trigonometric polynomials, shown in Figure 4, which approximates the flow for three cardiac cycles [30]. The length of three cycles (~ 3.3 seconds) allows for a total of 52 velocity data per measurement location using the sampling period Δ*t*_2_, and so the output space of <u>*g*</u> has dimension 52*N*. The outflow condition is a fully-absorbing boundary condition, described in more detail in [29].

**Figure 4:**
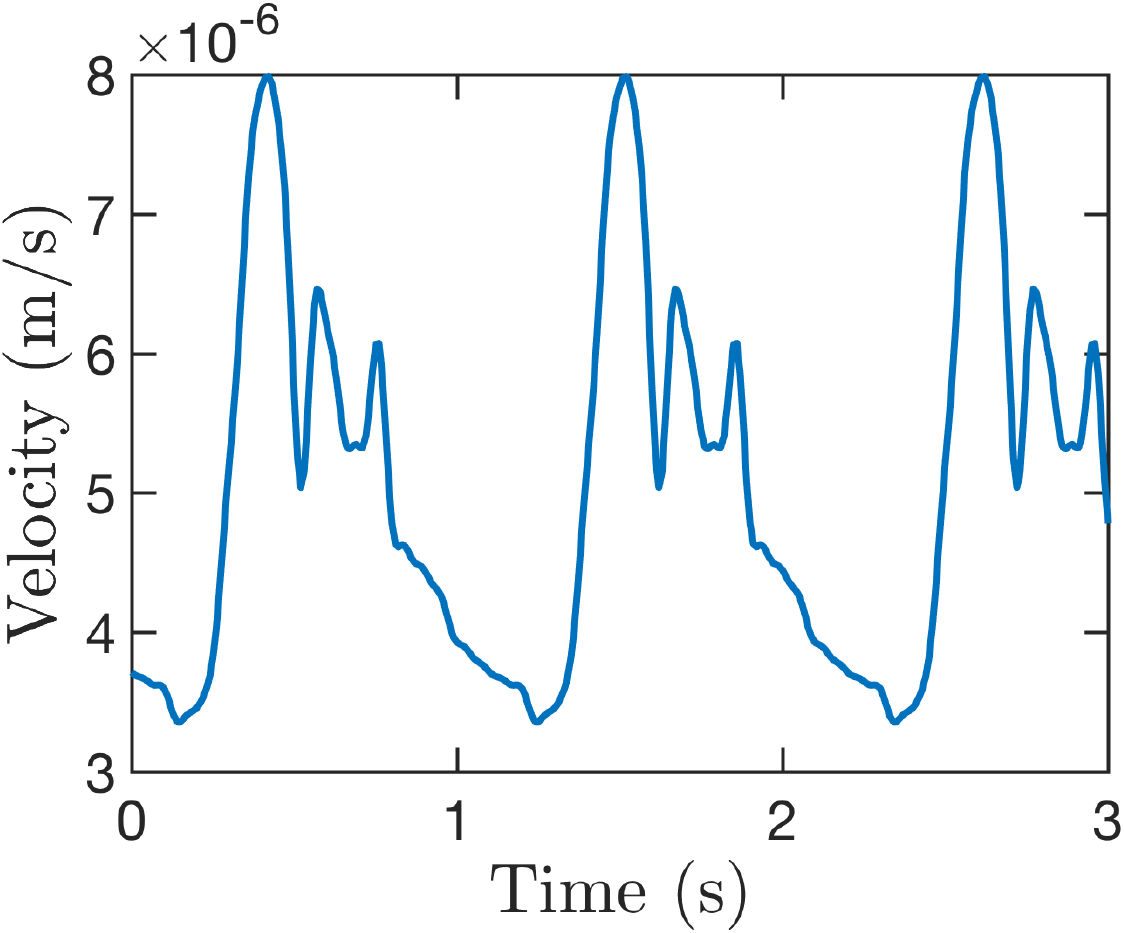
Inflow boundary condition for blood velocity *(m/s)* corresponding to three cardiac cycles.

We consider three free parameters: the blood viscosity *μ*, the arterial stiffness parameter *B*, and the relaxed cross-sectional area *A*_0_ (the last two of which vary by artery). A structural defect (e.g., an aneurysm or stenosis) can be modeled by varying the stiffness or relaxed area of a particular artery; to better emphasize the degree to which a flawed artery has been modified by its defect, results will use the scaled stiffness *β* = *B/B*\* (with respect to the reference stiffness *B*\*) and scaled cross-sectional area 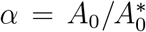 (with respect to the reference area 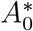), and so arteries with no defect will have *β* = *α* = 1.

We use our implementation of Bayesian uncertainty quantification to examine a number of questions in the context of this forward model, focusing in particular on the ability of uncertainty quantification to identify the location of structural flaws within the network using only noisy measurements of the flow velocity. To test the effectiveness of our implementation in these experiments thus requires noisy data <u>*D*</u> corresponding to a known truth; we use here synthetic data generated from the same model but with known, fixed parameters. It should be stressed that the approach is easily modified to admit real data and that there exist multiple practical methods for measuring blood flow velocities from in *vivo* arteries [36, 28, 35, 26]. Section 4.3 will show that flaws can be located accurately even when the parameters used to generate the synthetic data are significantly perturbed from the parameters used to perform uncertainty quantification.

Explicitly, observed data <u>*D*</u> are generated as

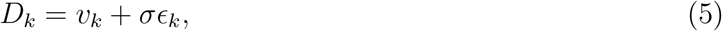

where *D_k_* is the noisy observation at time *t*_*k*_, *v_k_* is the flow velocity at time *t_k_*, *ϵ_k_* is a zero-mean, unit-variance Gaussian random variable, and *σ* is the noise level. Here, we choose *σ* to be a fraction *σ* = 0.01*η* (or sometimes 0.05*η*) of the standard deviation *η* of all velocity data *v_k_*.

In the following results, we use our implementation of uncertainty quantification to generate 500 samples from the posterior distribution *p*(*<u>θ</u>*|*<u>D</u>*, *M*) in a variety of scenarios. Posterior distributions are used for parameter estimation (Section 4.1) and to identify structural flaws via Bayesian model selection (Sections 4.2 and 4.3). Recovered posterior means, denoted with a hat (e.g., 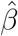), are used as parameter estimates in our analysis.

### 4.1 Parameter Estimation

We first consider a basic case of parameter estimation to illustrate the feasibility of the approach. Specifically, we estimate the blood viscosity *μ* and the scaled stiffness *β*_2_ of artery 2 (see Figure 3 for artery labels) assuming all other parameters are fixed to their reference values. The noisy data used, corrupted according to (5) with noise level *σ* = 0.01*η*, are sampled from a single location at the start of artery 6. As described in Section 3, we choose *σ* as an additional free parameter, requiring the approach to recover the noise level in addition to the target model parameters. A uniform distribution on [0.5,1.5] × [0.5,1.5] × [0,1] in the parameter space (*μ*, *β*_2_, *σ*) is used as the parameter prior *π*.

To determine the effect of the choice of sampling location, we additionally consider separate cases using data obtained from the start of arteries 1 and 8; for notational clarity, we refer to as Oi the case of observing the upflow end of artery *i*.

The results for the case *O*_6_ appear in Figure 5. *μ* and *β*_2_ are positively correlated in the posterior, i.e., simultaneously raising or lowering both the blood viscosity and the stiffness of artery 2 yields qualitatively similar observed data. Intuitively, in order to maintain a consistent rate of flow, a viscous flow necessitates more rigid artery walls.

**Figure 5:**
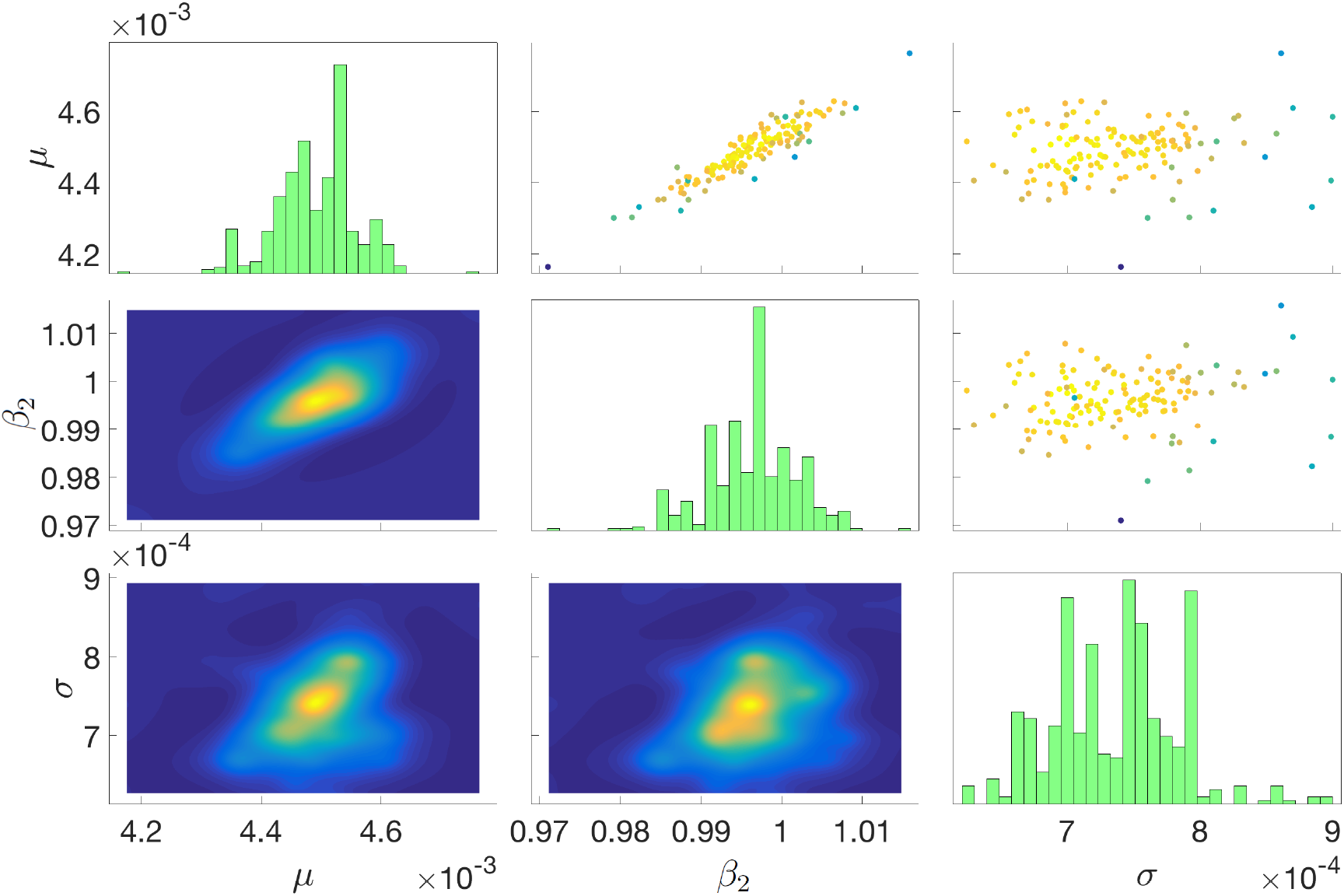
Parameter estimation results for blood viscosity *μ*, arterial stiffness *β*_2_ for artery 2, and noise level *σ* using corrupted reference data from the beginning of artery 6 (*O*_6_). Figures on the diagonal show histograms for each parameter. Subfigures below the diagonal show the marginal joint densities for each pair of parameters, while subfigures above the diagonal show the samples used in the final (convergent) stage of TMCMC. Colors correspond to likelihoods, with yellow likely and blue unlikely.

Numerical results for *O*_1_, *O*_6_, and *O*_8_ are summarized in Table 1. The recovered posterior means of (*μ*, *β*_2_, *σ*) were (0.00410, 0.972, 0.00074), (0.00449, 0.997, 0.00074), and (0.00447, 0.994, 0.00077), respectively, closely matching the true values *μ*\* = 0.0045 and 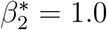 (*σ*\* differed by experiment due to differences in flow velocity by location: 0.00074, 0.00074, and 0.00077 for *O*_1_, *O*_6_, and *O*_8_, respectively). To quantify the degree of uncertainty in each parameter’s posterior distribution, we compute a coefficient of variation, defined as the ratio of the single-parameter posterior’s standard deviation to its mean (denoting the results 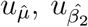, and 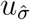); here, *O*_6_ and *O*_8_ recover parameters with comparatively lower uncertainty than *O*_1_, whose measurements were largely dominated by the inflow boundary condition. Nonetheless, in all cases the reference values used to generate the synthetic data were within one standard deviation of the recovered posterior means.

**Table 1:**
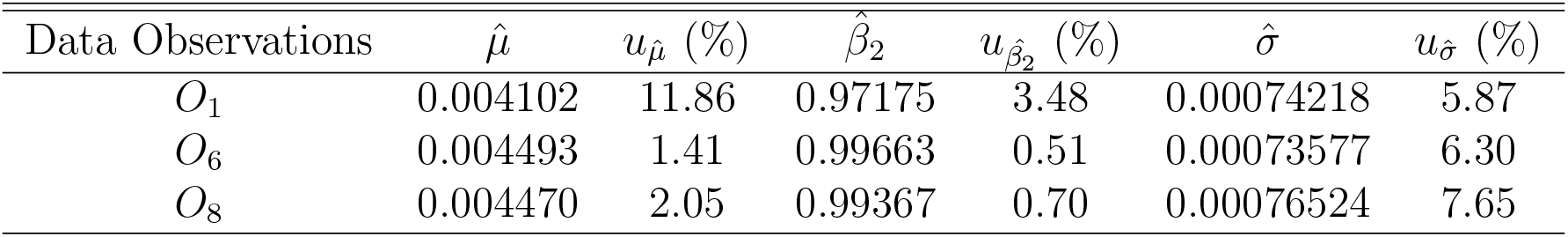
Posterior means and uncertainties for parameter estimation on the 19-artery network for three cases *O_i_* (noise level *σ* = 0.01*η*).

### 4.2 Locating Structural Flaws with Model Selection

Given the practicality of parameter estimation and its intermediate estimation of the model evidence *ρ*(*<u>D</u>*|*M*), the Bayesian model selection framework described in Section 3.2 is a natural approach to locating structural flaws in the arterial network. Namely, define as *M_i_* the model in which the scaled stiffness *β* of artery *i* has been perturbed from its reference value by an unknown amount, corresponding to, e.g., an aneurysm or stenosis. Parameter estimation as in Section 4.1 can be used to recover the perturbed stiffness which best matches the observed data, simultaneously yielding an estimate for the evidence *ρ*(*<u>D</u>*|*M_i_*) of model *M_i_*. Letting 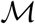 be the collection of *M_i_* for various arteries *i* in the network, the model selection distribution *Pr*(*M_i_*|*<u>D</u>*) is a probabilistic measure of the likelihood of the structural defect occurring in the artery *i* (as opposed to a different artery *j*). The class 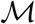 can easily be augmented with additional models; here, we also consider a model *M_i:j_* which freely varies the stiffness of two arteries *i* and *j*.

We first consider generating data *<u>D</u>* from a reference model using *β*_6_ = 0.5 and *β_i_* = 1, *i* ≠ 6, i.e., a model in which the scaled stiffness of artery 6 has been halved from its reference value. we consider three cases for data collection: a two sensor configuration using data from the end of arteries 1 and 7, a three sensor configuration using data from the end of arteries 1, 7, and 13, and a four sensor configuration using velocity data from the end of arteries 1, 7, 10, and 13. In each case, no sampling locations are adjacent to the damaged artery. The noise level is again chosen as *σ* = 0.01*η*, i.e., 1% Gaussian noise, and we employ the same uniform prior on [0, 3] × [0,1] for the parameters (*β, σ*).

Table 2 presents numerical results for six flaw models *M*_3_, *M*_6_, *M*_7_, *M*_11_, *M*_13_, and *M*_6:7_ when taking flow measurements from the ends of arteries 1 and 7. Models M_6_ and *M*_6:7_, both of which include the correct defect location in artery 6, are assigned the largest probabilities under the model selection posterior (*Pr*(*M_j_*|*<u>D</u>*) = 0.99949,0.00042, respectively); recovered parameter estimates 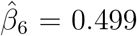 (*M*_6_) and 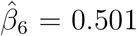 (*M*_6:7_) for the wall stiffness of the damaged artery were accurate to within one standard deviation. Though *M*_6:7_ assumes a second defect in artery 7, the posterior mean estimated the stiffness to be similar to the reference value 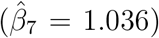. *M*_3_, *M*_7_, *M*_11_, and *M*_13_ are not able to accurately match the observed data, and so require a significantly higher noise level *σ* to explain differences between the evaluated and observed velocities. For this reason, these models are assigned negligible mass by *Pr*(*M_i_*|*<u>D</u>*).

**Table 2:** Numerical results for identification of a *β*_6_ = 0.5 aneurysm using noisy data from the ends of arteries 1 and 7 (noise level *σ* = 0.01*η*).

| Prediction Model | $\hat{\beta}$ | $u_{\hat{\beta}}$ (%) | $\hat{\sigma}$ | $u_{\hat{\sigma}}$ (%) | Log evidence | $Pr(M_j\vert \underline{D})$ |
| --- | --- | --- | --- | --- | --- | --- |
| $M_3$ | 0.741 | 2.82 | 0.000786 | 6.31 | 583.2 | 0.00009 |
| $M_6$ | 0.499 | 6.61 | 0.000729 | 3.73 | 592.5 | 0.99949 |
| $M_7$ | 1.055 | 3.14 | 0.000923 | 4.27 | 567.7 | $\sim 0$ |
| $M_{11}$ | 1.022 | 0.76 | 0.000892 | 5.96 | 566.3 | $\sim 0$ |
| $M_{13}$ | 0.587 | 5.81 | 0.000867 | 5.04 | 578.3 | $\sim 0$ |
| $M_{6:7}$ | [0.501, 1.036] | [3.75, 2.15] | 0.000719 | 5.16 | 584.7 | 0.00042 |

Note that the model *M*_6:7_ contains *M*_6_ in the sense that it can predict any combination of parameter values which *M*_6_ can predict. In this light, the relatively higher evidence for *M*_6_ over the broader error model M6:7 is in keeping with theoretical results available for Bayesian model class selection wherein over-parameterized model classes are penalized due to Occam’s factor [4].

Table 3 illustrates the corresponding results for a three-sensor configuration using blood flow velocity data from the ends of arteries 1, 7, and 13. The additional data collected from artery 13 significantly reduce the (already small) probabilities assigned to models other than *M*_6_ and *M*_6:7_. Interestingly, leveraging information from the end of artery 13, which is in a parallel tree (rather than directly upstream or downstream) from the damaged artery, has the effect of shifting mass from *M*_6_ to *M*_6:7_ in the model posterior, finding *Pr*(*M*_6_|*<u>D</u>*) = 0.708 and *Pr*(*M*_6:7_|*<u>D</u>*) = 0.292. Nonetheless, both *M*_6_ and *M*_6:7_ estimate the damaged stiffness *β*_6_ accurately (0.501 and 0.505, respectively), with *M*_6:7_ again finding the stiffness *β*_7_ of the undamaged artery 7 to be largely unchanged (1.031).

**Table 3:** Numerical results for identification of a *β*_6_ = 0.5 aneurysm using data from the ends of arteries 1, 7, and 13 (noise level *σ* = 0.01*η*).

| Prediction Model | $\hat{\beta}$ | $u_{\hat{\beta}}$ (%) | $\hat{\sigma}$ | $u_{\hat{\sigma}}$ (%) | Log evidence | $Pr(M_j\vert \underline{D})$ |
| --- | --- | --- | --- | --- | --- | --- |
| $M_3$ | 0.722 | 2.29 | 0.000811 | 4.81 | 875.6 | 0.00001 |
| $M_6$ | 0.501 | 4.31 | 0.000770 | 5.23 | 886.3 | 0.70756 |
| $M_7$ | 1.056 | 2.19 | 0.000956 | 4.64 | 853.1 | $\sim 0$ |
| $M_{11}$ | 1.005 | 0.60 | 0.000949 | 5.02 | 848.8 | $\sim 0$ |
| $M_{13}$ | 0.901 | 1.91 | 0.000887 | 5.06 | 858.3 | $\sim 0$ |
| $M_{6:7}$ | [0.505, 1.031] | [3.45, 2.33] | 0.000770 | 3.58 | 885.5 | 0.29242 |

Finally, Table 4 shows numerical results when velocity data are sampled at four monitoring locations: at the ends of arteries 1, 7, 10, and 13. The additional data from the end of artery 10 drive the model probabilities assigned to *M*_3_, *M*_7_, *M*_11_, and *M*_13_ down further (< 10^−8^), rendering them orders of magnitude smaller than the probabilities assigned to M6 and M6:7 (0.9996 and 0.0004, respectively). The estimated scaled stiffness remains accurate to within one standard deviation, with *M*_6_ and *M*_6:7_ finding 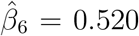 and 0.521, respectively, and *M*_6:7_ again estimates the stiffness of artery 7 to be only slightly perturbed 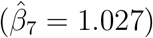.

**Table 4:** Numerical results for identification of a *β*_6_ = 0.5 aneurysm using data from the ends of arteries 1, 7, 10, and 13 (noise level *σ* = 0.01*η*).

| Prediction Model | $\hat{\beta}$ | $u_{\hat{\beta}}$ (%) | $\hat{\sigma}$ | $u_{\hat{\sigma}}$ (%) | Log evidence | $Pr(M_j\vert \underline{D})$ |
| --- | --- | --- | --- | --- | --- | --- |
| $M_3$ | 0.760 | 2.60 | 0.000817 | 3.93 | 1169.9 | $\sim 0$ |
| $M_6$ | 0.520 | 2.57 | 0.000762 | 3.48 | 1187.3 | 0.9996 |
| $M_7$ | 1.046 | 2.72 | 0.000926 | 3.86 | 1144.5 | $\sim 0$ |
| $M_{11}$ | 0.996 | 0.19 | 0.000932 | 3.58 | 1134.1 | $\sim 0$ |
| $M_{13}$ | 0.890 | 1.83 | 0.000902 | 3.47 | 1148.8 | $\sim 0$ |
| $M_{6:7}$ | [0.521, 1.027] | [3.98, 2.49] | 0.000771 | 3.63 | 1179.5 | 0.0004 |

Taken together, these configurations support two conclusions about Bayesian model selection for flaw identification: first, that increasing the number of locations at which data are sampled reduces the probabilities assigned to incorrect models, and second, that model selection can accurately determine the defect location and magnitude for a variety of sensor configurations, including configurations which do not sample from at or near the defect location.

#### 4.2.1 Model Selection for Cross-Sectional Area

As previously suggested, aneurysms and stenoses can also be modeled by adjusting the initial crosssectional area of an artery rather than its stiffness. Ideally, the Bayesian framework for model selection should provide similar results when the stiffnesses *β* are fixed and models Mi instead allow the scaled cross-sectional area *α* of the defective artery to be perturbed. In what follows, we examine similar scenarios to the above in the case where, rather than reducing its wall stiffness, the relaxed cross-sectional area of artery 6 is altered. A uniform prior on [0, 3] × [0, 1] is used for the parameters (*α, σ*).

We first consider the case *α*_6_ = 1.5, i.e., an aneurysm in which the defective artery (again, artery 6 in the reference model) has become enlarged by 50%. Noisy flow velocity data are collected from the ends of arteries 1, 7, 10, and 13, as in the final case of the previous section; results appear in Table 5. *M*_6_ and *M*_6:7_ are again the most likely models (*Pr*(*M_j_*|*<u>D</u>*) = 0.998 and 0.002, respectively), suggesting that the previous results do not rely on the specific choice of the parameter *β*. Other models were assigned negligible probabilities. Similarly to the results for reduced stiffness, both M6 and *M*_6:7_ accurately recover the defect magnitude (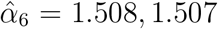, respectively), and *M*_6:7_ finds artery 7 to be unchanged 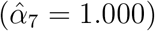.

**Table 5:** Numerical results for area-based identification of an *α*_6_ = 1.5 aneurysm using data from the ends of arteries 1, 7, 10, and 13 (noise level *σ* = 0.01*η*).

| Prediction Model | $\hat{\alpha}$ | $u_{\hat{\alpha}}$ (%) | $\hat{\sigma}$ | $u_{\hat{\sigma}}$ (%) | Log evidence | $Pr(M_j\vert \underline{D})$ |
| --- | --- | --- | --- | --- | --- | --- |
| $M_3$ | 1.126 | 26.29 | 0.00554 | 3.40 | 781.0 | $\sim 0$ |
| $M_6$ | 1.508 | 0.45 | 0.00078 | 4.33 | 1183.6 | 0.998 |
| $M_7$ | 0.976 | 0.17 | 0.00415 | 4.15 | 827.9 | $\sim 0$ |
| $M_{11}$ | 1.048 | 0.48 | 0.00471 | 3.41 | 810.0 | $\sim 0$ |
| $M_{13}$ | 1.014 | 0.17 | 0.00518 | 3.18 | 793.0 | $\sim 0$ |
| $M_{6:7}$ | [1.507, 1.000] | [0.52, 0.031] | 0.00077 | 3.24 | 1177.6 | 0.002 |

We then consider the same scenario for a reduction *α*_6_ = 0.5 in the cross-sectional area of artery 6, i.e., a stenosis in which the defective artery has narrowed by 50%. Results are summarized in Table 6. *M*_6_ and *M*_6:7_ recover the reduced area accurately (*α*_6_ = 0.500,0.501, respectively) and are assigned the highest model evidence (*Pr*(*M_j_*|<u>*D*</u>) ≈ 1.00 and ~ 10^−4^, respectively).

**Table 6:** Numerical results for area-based identification of an *α*_6_ = 0.5 stenosis using data from the ends of arteries 1, 7, 10, and 13 (noise level *σ* = 0.01*η*).

| Prediction Model | $\hat{\alpha}$ | $u_{\hat{\alpha}}$ (%) | $\hat{\sigma}$ | $u_{\hat{\sigma}}$ (%) | Log evidence | $Pr(M_j\vert \underline{D})$ |
| --- | --- | --- | --- | --- | --- | --- |
| $M_3$ | 0.072 | 8.00 | 0.023 | 4.04 | 478.9 | $\sim 0$ |
| $M_6$ | 0.500 | 0.05 | 0.00076 | 3.16 | 1178.0 | 1.00 |
| $M_7$ | 1.127 | 0.78 | 0.0195 | 4.83 | 513.3 | $\sim 0$ |
| $M_{11}$ | 0.790 | 2.37 | 0.0215 | 3.72 | 493.8 | $\sim 0$ |
| $M_{13}$ | 0.941 | 0.73 | 0.0238 | 3.33 | 475.2 | $\sim 0$ |
| $M_{6:7}$ | [0.501, 1.000] | [0.08, 0.03] | 0.00076 | 3.30 | 1164.6 | $\sim 0$ |

Table 7 shows results for the same magnitude stenosis (*α*_6_ = 0.5) with increased observational noise level *σ* = 0.05*η*. The log evidence of models *M*_6_ and *M*_6:7_ is sharply reduced compared to Table 6, though M6 and M6:7 remain the most probable models under the model selection posterior, with *Pr*(*M_j_*|*<u>D</u>*) = 0.983 and 0.017, respectively. Both models additionally recover the reduced area accurately (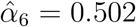 and 0.503, respectively) despite the increased noise.

**Table 7:**
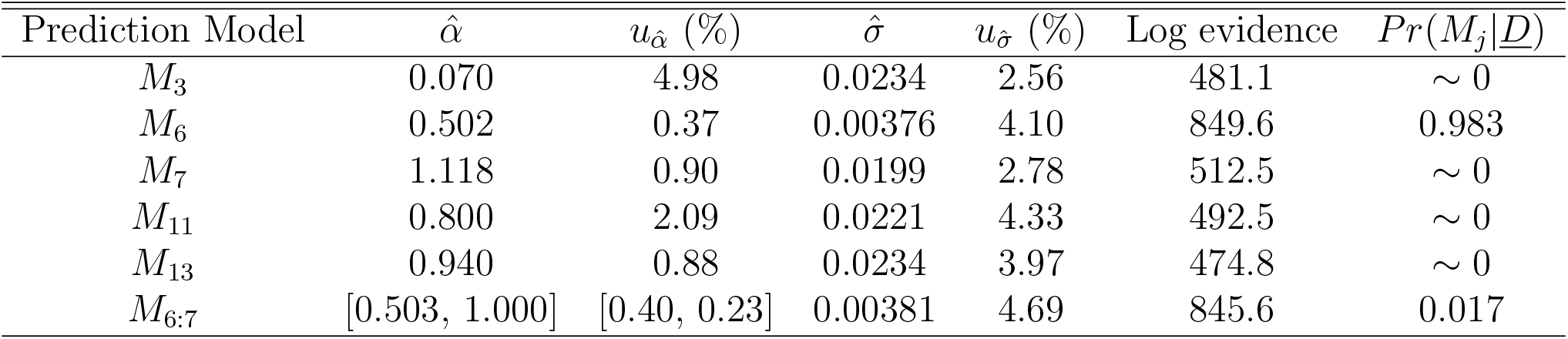
Numerical results for area-based identification of an *α*_6_ = 0.5 stenosis using data from the ends of arteries 1, 7, 10, and 13 (noise level *σ* = 0.05*η*).

Finally, Table 8 considers the case of a smaller-magnitude stenosis (*α*_6_ = 0.8). Results were similar to those of Table 7, with accurate recovery of location *(Pr(Mj*|<u>*D*</u>) = 0.99999 and 0.00001 for Me, *M*_6:7_, respectively) and magnitude (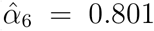 and 0.802). As in the previous area-modification scenarios, *M*_6:7_ found artery 7 to be unaffected 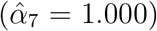, thereby coinciding with the single defect model *M*_6_.

**Table 8:** Numerical results for area-based identification of an *α_6_* = 0.8 stenosis using data from the ends of arteries 1, 7, 10, and 13 (noise level *σ* = 0.01*η*).

| Prediction Model | $\hat{\alpha}$ | $u_{\hat{\alpha}}$ (%) | $\hat{\sigma}$ | $u_{\hat{\sigma}}$ (%) | Log evidence | $Pr(M_j\vert \underline{D})$ |
| --- | --- | --- | --- | --- | --- | --- |
| $M_3$ | 0.175 | 4.13 | 0.00493 | 3.71 | 809.0 | $\sim 0$ |
| $M_6$ | 0.801 | 0.18 | 0.00075 | 4.68 | 1185.1 | 0.99999 |
| $M_7$ | 1.024 | 0.15 | 0.00404 | 3.78 | 838.0 | $\sim 0$ |
| $M_{11}$ | 0.955 | 0.45 | 0.00447 | 3.92 | 820.5 | $\sim 0$ |
| $M_{13}$ | 0.987 | 0.17 | 0.00488 | 3.89 | 802.7 | $\sim 0$ |
| $M_{6:7}$ | [0.802, 1.000] | [0.16, 0.033] | 0.00077 | 3.11 | 1173.7 | 0.00001 |

### 4.3 Locating Defects with a Misspecified Model

Results have so far assumed the model selection framework is provided the reference values for all model parameters, i.e., the non-defective stiffness and area of each artery are known. In a scenario using real-world data, these “known” values may themselves be estimated from noisy measurements. A final but crucial test of the robustness of the model selection flaw identification framework is thus to perform experiments in which the reference parameters used by the method are incorrect, and so no combination of free parameters is capable of reproducing the observed data.

We now revisit the cases of Section 4.2.1, beginning with the case of an *α*_6_ = 1.5 aneurysm in artery 6. In addition to corrupting observed flow velocities with additive Gaussian noise, we now additionally corrupt the parameters themselves: the initial cross-sectional area *α_k_* for each artery *k* is noised as

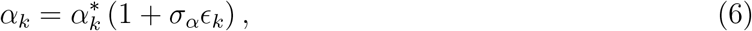

where 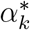 is the reference value, *ϵ_k_* is again a standard normal random variable, and *σ_α_* is the parameter noise level. The structural parameters used to generate the synthetic data (*α_k_* from Eq. 6) thus differ from the fixed values used in the defect models 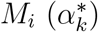.

As before, Bayesian model selection is performed assuming the prediction equation (3), which is now misspecified (it assumes correctness of the reference parameters 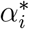). As a result, the 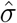 estimated by posterior samples must now capture the effects of both the true observational noise level *σ* and the parameter noise level *σ_α_*.

Table 9 shows numerical results for Bayesian model selection in this setting. Despite the mis-specification, *M*_6_ and *M*_6:7_ again dominate the model posterior, with *Pr*(*M*_6_|*<u>D</u>*) = 0.907 and *Pr*(*M*_6:7_|*<u>D</u>*) = 0.093, respectively. Both overestimate the defect magnitude (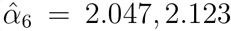, respectively), though *M*_6:7_ again estimates artery 7 to be unaffected 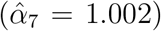. We note that some error in 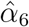 is expected, as it attempts to fit observations from the noised-parameter model and thus varies significantly depending on the particular values of *α_k_* from Eq. (6). Despite this effect, identification of the location appears robust to perturbation of model parameters, with all other models assigned negligible probability (*Pr*(*M_i_*|*<u>D</u>*) ~ 0).

**Table 9:** Numerical results for area-based identification of an *α*_6_ = 1. 5 aneurysm using data from the ends of arteries 1, 7, 10, and 13 (noise level *σ* = 0.01*η*) with misspecified cross-sectional areas (perturbed with noise level *σ_α_* = 0.01).

| Prediction Model | $\hat{\alpha}$ | $u_{\hat{\alpha}}$ (%) | $\hat{\sigma}$ | $u_{\hat{\sigma}}$ (%) | Log evidence | $Pr(M_j\vert \underline{D})$ |
| --- | --- | --- | --- | --- | --- | --- |
| $M_3$ | 1.312 | 29.2 | 0.00761 | 3.39 | 711.6 | $\sim 0$ |
| $M_6$ | 2.047 | 0.89 | 0.00217 | 3.98 | 972.9 | 0.907 |
| $M_7$ | 0.970 | 0.23 | 0.00628 | 3.48 | 748.7 | $\sim 0$ |
| $M_{11}$ | 1.075 | 0.56 | 0.00645 | 3.76 | 746.9 | $\sim 0$ |
| $M_{13}$ | 1.016 | 0.29 | 0.00725 | 3.53 | 716.8 | $\sim 0$ |
| $M_{6:7}$ | [2.123, 1.002] | [3.27, 0.11] | 0.00216 | 4.32 | 970.6 | 0.093 |

Turning to the second case (*α*_6_ = 0.5), model selection again successfully locates the defect despite the misspecification (Table 10), with *M*_6_ assigned nearly all mass by the model selection posterior. In this case, parameter estimation recovers the defect magnitude accurately 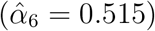. In keeping with previous results, defect model M6:7 finds a similar reduction in cross-sectional area for the damaged artery 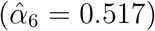 and little change in the defect-free artery 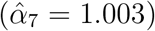.

**Table 10:** Numerical results for area-based identification of an *α*_6_ = 0.5 stenosis using data from the ends of arteries 1, 7, 10, and 13 (noise level *σ* = 0.01*η*) with misspecified cross-sectional areas (perturbed with noise level *σ_α_* = 0.01).

| Prediction Model | $\hat{\alpha}$ | $u_{\hat{\alpha}}$ (%) | $\hat{\sigma}$ | $u_{\hat{\sigma}}$ (%) | Log evidence | $Pr(M_j\vert \underline{D})$ |
| --- | --- | --- | --- | --- | --- | --- |
| $M_3$ | 0.816 | 7.29 | 0.0220 | 4.36 | 485.7 | $\sim 0$ |
| $M_6$ | 0.515 | 0.17 | 0.0021 | 3.22 | 973.6 | 0.999 |
| $M_7$ | 1.118 | 0.66 | 0.0181 | 3.44 | 529.0 | $\sim 0$ |
| $M_{11}$ | 0.808 | 1.72 | 0.0207 | 3.41 | 505.7 | $\sim 0$ |
| $M_{13}$ | 0.943 | 0.69 | 0.0221 | 3.14 | 490.5 | $\sim 0$ |
| $M_{6:7}$ | [0.517, 1.003] | [0.23, 0.001] | 0.0021 | 3.87 | 966.5 | 0.001 |

The third case repeated the *α*_6_ = 0.5 experiment with increased observational noise *σ* = 0.05*η*; results for the same case with parameter noise (now also increased to *σ*_*α*_ = 0.05) are shown in Table 11. *M*_6:7_ is significantly more likely than in previous cases (*Pr*(*M*_6:7_|*<u>D</u>*) = 0.783), though *M*_6_ is still assigned all remaining posterior mass (*Pr*(*M*_6_|*<u>D</u>*) = 0.217). The recovered uncertainties 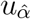 are significantly higher than in Table 10 due to the higher level of noise, with both *M*_6_ and *M*_6:7_ underestimating the magnitude of the damage (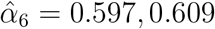, respectively).

**Table 11:** Numerical results for area-based identification of an *α*_6_ = 0.5 stenosis using data from the ends of arteries 1, 7, 10, and 13 (noise level *σ* = 0.05*η*) with misspecified model parameters (perturbed with noise level *σ_α_* = 0.05).

| Prediction Model | $\hat{\alpha}$ | $u_{\hat{\alpha}}$ (%) | $\hat{\sigma}$ | $u_{\hat{\sigma}}$ (%) | Log evidence | $Pr(M_j\vert \underline{D})$ |
| --- | --- | --- | --- | --- | --- | --- |
| $M_3$ | 1.668 | 45.9 | 0.0195 | 4.35 | 518.5 | $\sim 0$ |
| $M_6$ | 0.597 | 0.80 | 0.0111 | 3.31 | 636.6 | 0.217 |
| $M_7$ | 1.085 | 0.62 | 0.00076 | 4.82 | 558.5 | $\sim 0$ |
| $M_{11}$ | 0.894 | 1.52 | 0.0185 | 3.82 | 527.1 | $\sim 0$ |
| $M_{13}$ | 0.947 | 0.51 | 0.0175 | 3.62 | 534.9 | $\sim 0$ |
| $M_{6:7}$ | [0.609, 1.015] | [1.22, 0.44] | 0.0108 | 2.22 | 637.9 | 0.783 |

Finally, Table 12 shows results for the fourth case (*α*_6_ = 0.8) in the presence of *σ_α_* = 0.01 parameter noise. While model selection again recovers the correct defect location (*Pr*(*M_j_*|*<u>D</u>*) = 0.969, 0.031 for *M*_6_, *M*_6:7_, respectively), the smaller-magnitude stenosis proves more challenging for parameter estimation, with 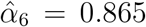 and 0.869, respectively, notably underestimating the magnitude of the damage.

**Table 12:** Numerical results for area-based identification of an *α*_6_ = 0.8 stenosis using data from the ends of arteries 1, 7, 10, and 13 (noise level *σ* = 0.01*η*) with misspecified model parameters (perturbed with noise level *σ_α_* = 0.01).

| Prediction Model | $\hat{\alpha}$ | $u_{\hat{\alpha}}$ (%) | $\hat{\sigma}$ | $u_{\hat{\sigma}}$ (%) | Log evidence | $Pr(M_j\vert \underline{D})$ |
| --- | --- | --- | --- | --- | --- | --- |
| $M_3$ | 1.767 | 22.2 | 0.00388 | 3.57 | 852.4 | $\sim 0$ |
| $M_6$ | 0.865 | 0.35 | 0.00204 | 3.36 | 974.9 | 0.969 |
| $M_7$ | 1.017 | 0.13 | 0.00320 | 3.87 | 889.5 | $\sim 0$ |
| $M_{11}$ | 0.979 | 0.24 | 0.00369 | 3.61 | 860.3 | $\sim 0$ |
| $M_{13}$ | 0.988 | 0.12 | 0.00345 | 3.49 | 869.5 | $\sim 0$ |
| $M_{6:7}$ | [0.869, 1.003] | [0.50, 0.076] | 0.00209 | 3.22 | 971.5 | 0.031 |

## 5 Discussion

Taken together, the results describe a robust approach for uncertainty quantification in the context of arterial networks. The model selection posterior universally assigned the highest probabilities (by several orders of magnitude) only to those models which included the true defect location, even in cases where simulated data were sparse, noisy, and poorly located. The Bayesian uncertainty quantification framework thus appears a powerful tool for comparing and fitting models.

Though all results were generated using simulated noisy data, they simultaneously suggest that the approach would prove useful for real-world inference. The experiments outlined in Section 4.2 show the method to successfully recover parameter values (often within one standard deviation) and identify the defect location in a range of sampling cases which varied sensor numbers and locations, and so the approach is not reliant on a particular set of observed data which may not be realistically attainable. Results were also consistent when using alternative magnitudes and parametrizations of arterial defects (the scaled cross-sectional area *α* and boundary stiffness *β*) and using models which considered different numbers of defects (in particular, the two-defect model *M*_6:7_ which consistently found the “defective” artery 7 to be largely unaltered). Most importantly, Section 4.3 showed inference to remain effective even when the model used to generate the simulated data differed from the model used to perform inference (i.e., model misspecification). As mathematical models are inherently simplifications of complex physical systems, robustness to misspecification is an essential component of applicability to experimental data.

The approach itself readily facilitates the incorporation of real data, which can be used in place of simulated data without otherwise altering the method. A natural extension is thus direct application to medical datasets. Future work will also incorporate alternative network structures and models and will consider defects in more localized arterial subdomains.

## 6 Acknowledgements

Parts of this research were conducted using computational resources and services at the Center for Computation and Visualization, Brown University. KL and AM were partially supported by the NSF through grants DMS-1521266 and DMS-1552903. PK was supported by the European Research Council Advanced Investigator Award (Grant: 341117).

## Appendix A High-performance implementations

Π4U [15] is a platform-agnostic task-based UQ framework that supports nested parallelism and automatic load balancing in large scale computing architectures. The software is open-source and includes HPC implementations for both multicore and GPU clusters of algorithms such as Transitional Markov chain Monte Carlo (TMCMC) and Approximate Bayesian Computational Subset-simulation. The irregular, dynamic and multi-level task-based parallelism of the algorithms (Fig. 6, left) is expressed and fully exploited by means of the TORC runtime library [3]. TORC is a software library for programming and running unaltered task-parallel programs on both shared and distributed memory platforms. TORC orchestrates the scheduling of function evaluations on the cluster nodes (Fig. 6, right). The parallel framework includes multiple features, most prominently the inherent load balancing, fault-tolerance and high reusability. The TMCMC method within Π4U is able to achieve an overall parallel efficiency of more than 90% on 1024 compute nodes of Swiss supercomputer Piz Daint running hybrid MPI+GPU molecular simulation codes with highly variable time-to-solution between simulations with different interaction parameters.

**Figure 6:**
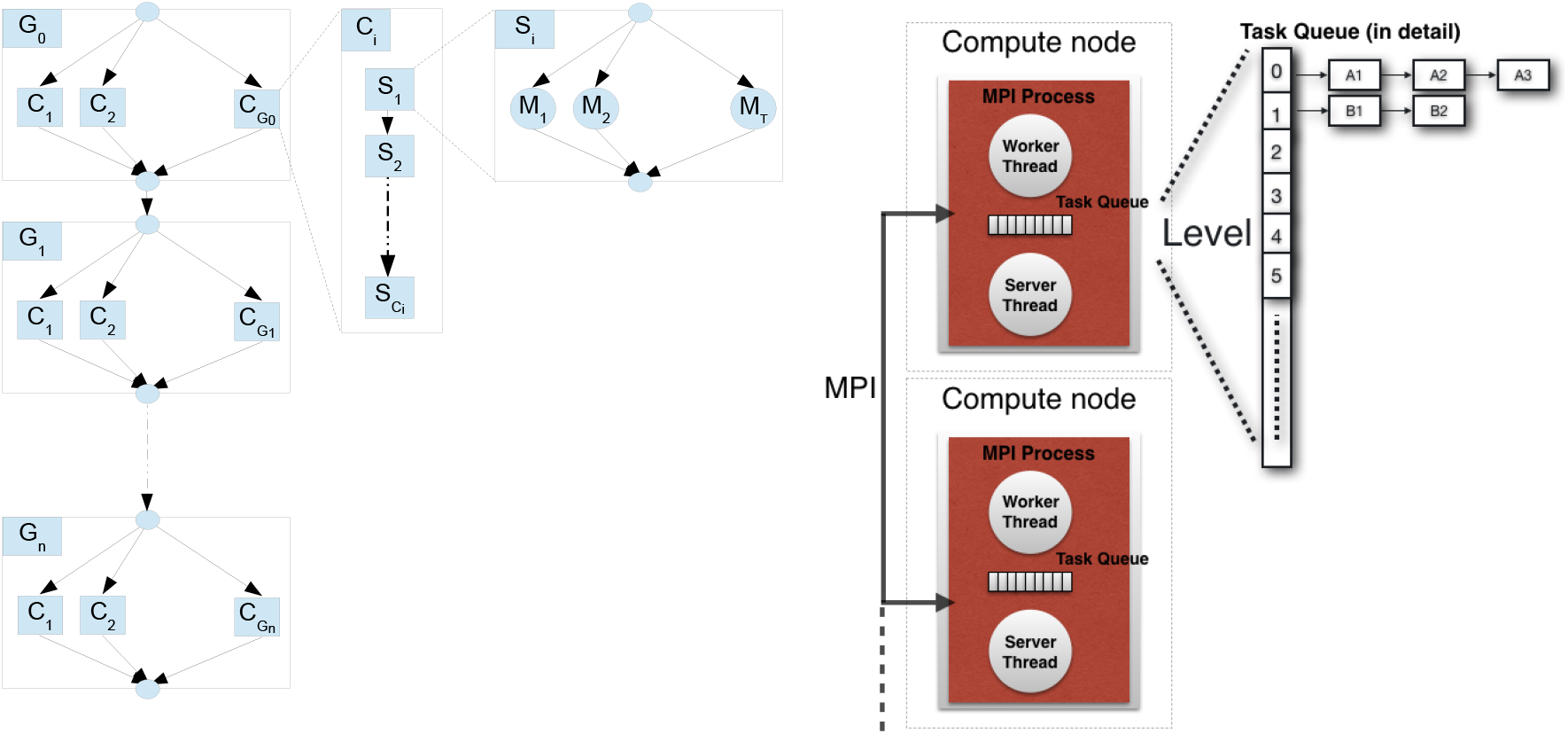
Task graph of the TMCMC algorithm (left) and parallel architecture of the TORC library (right).

